# Automatic structural analysis of bioinspired percolating network materials using graph theory

**DOI:** 10.1101/2021.04.07.438877

**Authors:** Drew Vecchio, Samuel Mahler, Mark D. Hammig, Nicholas A. Kotov

## Abstract

Mimicking numerous biological membranes and nanofiber-based tissues, there are multiple materials that are structured as percolating nanoscale networks (PPNs). They reveal unique combination of properties and the family of PNN-based composites and nanoporous materials is rapidly expanding. Their technological significance and the necessity of their structural design require a unifying approach for their structural description. However, their complex aperiodic architectures are difficult to describe using traditional methods that are tailored for crystals. A related problem is the lack of computational tools that enable one to capture and enumerate the patterns of stochastically branching fibrils that are typical for these composites. Here, we describe a conceptual methodology and a computational package, *StructuralGT,* to automatically produce a graph theoretical (GT) description of PNNs from various micrographs. Using nanoscale networks formed by aramid nanofibers (ANFs) as examples, we demonstrate structural analysis of PNNs with 13 GT parameters. Unlike qualitative assessments of physical features employed previously, *StructuralGT* allows quantitative description of the complex structural attributes of PNNs enumerating the network’s morphology, connectivity, and transfer patterns. Accurate conversion and analysis of micrographs is possible for various levels of noise, contrast, focus, and magnification while a dedicated graphical user interface provides accessibility and clarity. The GT parameters are expected to be correlated to material properties of PNNs (e.g. ion transport, conductivity, stiffness) and utilized by machine learning tools for effectual materials design.

**Table of Content:** 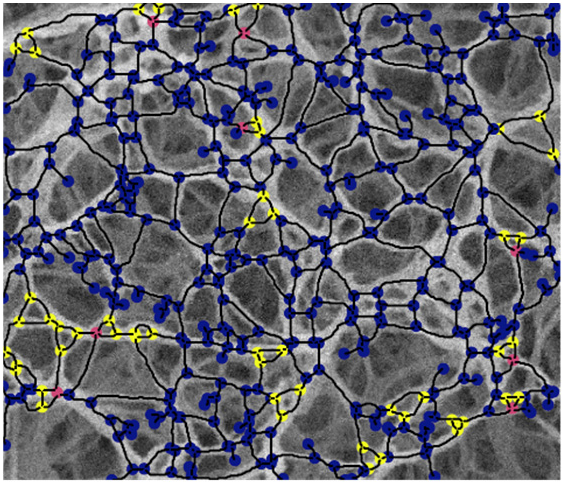

## 1. Introduction

Finding predictive relations between the microscale structure of a solid and its macroscopic properties is a long-standing problem in physics, chemistry, biology, and materials science. This task becomes even more challenging when the structure of the material does not reveal a periodic or quasiperiodic^[1]^ pattern typically used to establish the structure-property relationships. The problem of identifying the structural descriptors becomes particularly perplexing for composites with complex architectures that can be described as percolating nanoscale networks (PNNs, **Figure 1**). Such organization is common for numerous biological materials, for example cartilage, basal and glomerular membranes.^[2,3]^ Being self-assembled from a variety of peptides, amyloid fibers, for instance in bacterial biofilms,^[4]^ also display the similar percolating pattern. PNNs can also form self-assemble from multiple types of abiological nanofibers, nanoplatelets, and nanoparticles.^[5–19]^ This family of composite and porous materials is rapidly expanding due to the simplicity of their preparation as well as the increasing multiplicity of nanoscale components from which they can form. Architectures with similarly complex organization that possess variable degrees of stochasticity can also be found in some micro and macroscale materials, which accentuates the need for the development of suitable methods for their structural description. The hybrid and porous materials that possess a percolating network architecture formed from nanoscale components represents a harder and more impactful problem for structural description and materials design because: (**1**) PNNs encompass a large set of load-bearing composites, membranes, catalysts, biomaterials, thermal insulators, and energy materials. and (**2**) although the transition to periodic architectures is possible for micro- and macroscale lattices due to 3D printing technologies, that simplification is not possible for PNNs because they are typically formed via self-assembly, which is ubiquitous and energy efficient.^[5,6,12,13]^

**Figure 1.**
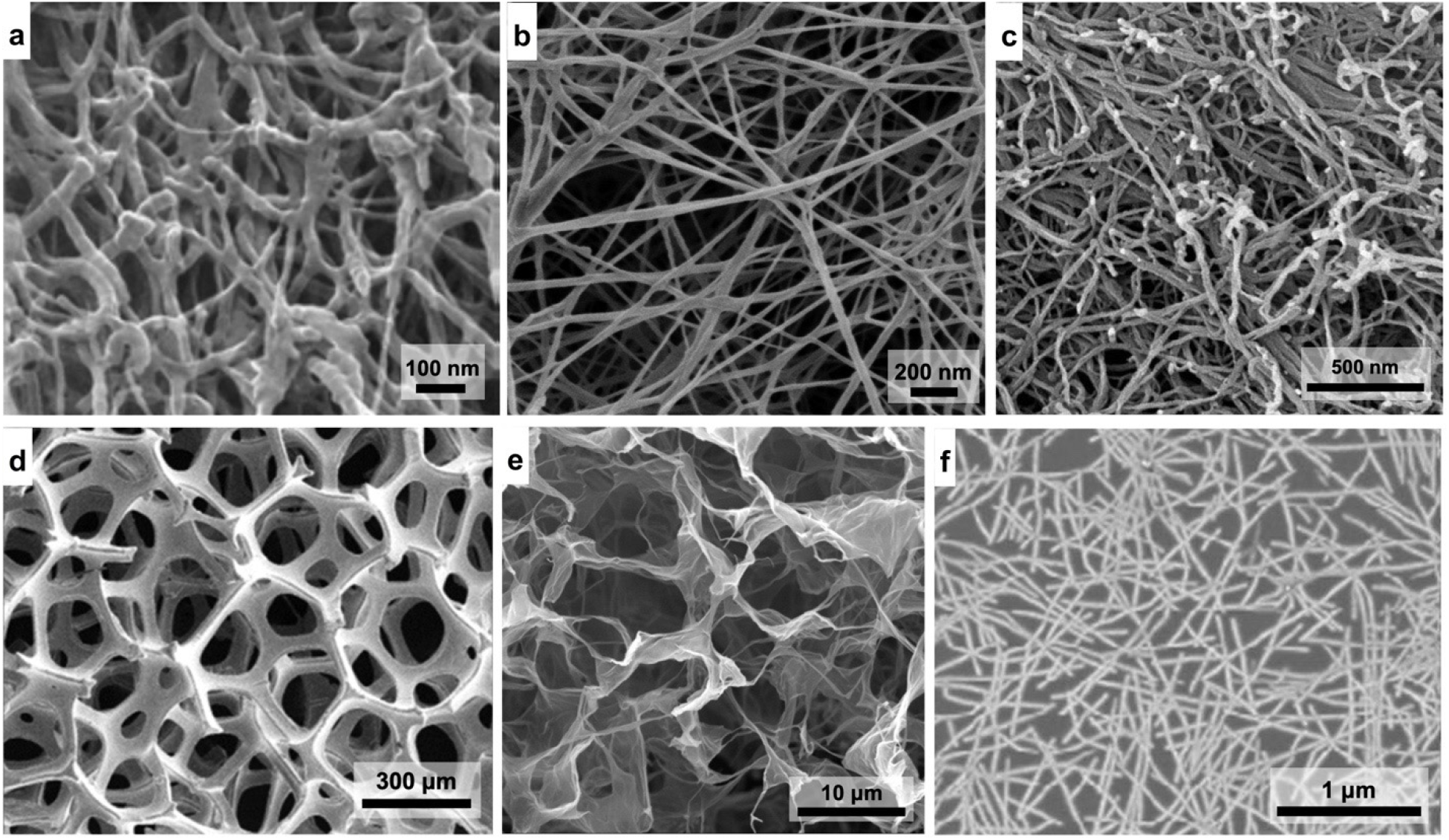
Images of several types of structural networks that can be successfully characterized by applying graph theory. **a)** Helium ion microscopy image of an articular cartilage collagen network. Reproduced with permission.^[15]^ 2012, John Wiley and Sons. **b)** SEM image of a freeze-dried bacterial cellulose network, with surfaces modified by trimethylsilyl groups. Reproduced with permission.^[16]^ 2019, American Chemical Society. **c)** SEM image of boron nitride nanotubes. Reproduced with permission.^[10]^ 2005, The Royal Society of Chemistry. **d)** SEM image of Ni foam with bulk density *ρ* = 0.16 g cm^−3^. Reproduced with permission.^[28]^ 2013, American Chemical Society. **e)** Hydrogel from self-assembled graphene oxide sheets with Ca^2+^ ions. Reproduced with permission.^[9]^ 2011, American Chemical Society. **f)**SEM image of a single walled carbon nanotube network on a bare SiO2 substrate. Reproduced with permission.^[19]^ 2015, Elsevier.

Some examples of PNNs include hybrid materials based on networks formed by silver nanowires,^[14]^ collagen fibers,^[15,16]^ nanofibrous cellulose,^[11,17,20]^ clay platelets,^[18]^ carbon nanotubes,^[6,13,19,21]^ aramid nanofibers (ANFs),^[7,8]^ graphene,^[9]^ boron nitride nanotubes,^[10]^ and others. Assessing the existing possibilities through which one can make quantitative descriptions of their structures at the scales from tens of nanometers to tens of micrometers, the metrics of fibril length, pore size/distribution, wall thickness, volume fraction, and defect sites can be applied.^[22–25]^ However, the properties of PNN materials cannot be comprehensively and uniquely determined by these structural parameters because this set of descriptors lacks the characterization of the material’s architecture as a whole, which is essential for conductivity, stiffness, transparency, and many other properties.^[26,27]^ A similar challenge also emerges in foams with a stochastic distribution of interconnected micropores made from polymers, ceramics, and metals,^[28]^ whose architectures are congruent to PNNs.^[29]^

The need for the comprehensive structural quantification of materials with aperiodic stochastic architectures inclusive of both ‘local’ and ‘global’ structural patterns becomes, thus, general and fundamental. A closely related problem is the lack of computational tools and algorithms to enumerate the materials with non-crystalline architecture that would be rapid and accurate. The need for such tools capable of processing large amount of micrographs similar to those in **Figure 1** from which descriptors can be extracted is further amplified by rapidly developing machine learning (ML) methods for materials science.^[30–35]^ Considering the current practices of image labeling and the capabilities of common image processing software,^[24,25,36,37]^ it is important to increase their throughput and minimize bias from experimentalists.

A systematic protocol for the description of materials with percolating networks can be accomplished based on graph theory (GT), also known as the network theory. A graph, *G(n, e)*, is a mathematical model consisting of data points (nodes, *n*) joined by lines (edges, *e*) that follow a specific protocol reflecting the constitutive or structural relationships between them. GT has long been used to describe financial, informational, social, and biological networks,^[38–43]^ but not the structure of nanoscale materials. The key challenge for using GT for the description of nanomaterials is the adequate translation of their physical structure into a *G(n, e)* model.^[44]^ Here we propose that PNNs can be generally represented by *G(n,e)* by describing continuous nanofilaments, nanofibers, or nanowires as edges while their intersections are identified as nodes. The resulting *G(n, e)* representation affords multiple and diverse measures to comprehensively describe virtually any type of PNN regardless of the dimensions of the constitutive components and complexity of the resulting network. Recent data also indicate that GT indexes can not only be directly correlated with some macroscale properties but they can also enumerate the structural differences between PNNs with seemingly identical architectures.^[44,45]^

It is theoretically possible to create *G(n, e)* models that can be defined as edge or adjacency matrixes for a variety of materials in a high-throughput manner by analyzing numerous electron microscopy images, but the computational tools and integrated mathematical algorithms for such a task are missing.^[46]^ To address this need we created an open-source Python software package, *StructuralGT* that takes as an input digital images obtained by various microscopy techniques, such as scanning electron microscopy (SEM) and transmission electron microscopy (TEM). The outputs include: (**1**) *G(n,e)* describing the material, (**2**) histograms of GT parameters, (**3**) their ‘heat maps’ overlaid upon the original electron microscopy images, and (**4**) their average values. *StructuralGT* employs a graphical user interface (GUI) to make the software accessible and streamlined, while also providing flexibility for the input image and desired GT measurements. The current structural descriptors include average nodal connectivity 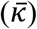, network diameter, graph density (*D*), global efficiency (*E*), average clustering coefficient (*Δ*), Wiener Index (*WI*), assortativity coefficient (*r*), betweenness centrality (*C_B_*), closeness centrality (*C_C_*), eigenvector centrality (*C_E_*), their weighted analogs, and other GT parameters describing both local and global organization of the nanofilament network.

## 2. Image and structural analysis of PNNs by *StructuralGT*

*StructuralGT* is freely available as either a Python package or a stand-alone executable file via GitHub (https://github.com/drewvecchio/StructuralGT). The workflow of *StructuralGT* is divided into three sections – (1) image processing, (2) graph extraction, and (3) the calculation of GT parameters (that can also be referred to as descriptors or indexes). In brief, the image processing segment converts the gray scale microscopy image into a binary image, from which the graph extraction module converts the latter into a dictionary-of-dictionaries-of-dictionaries Python data structure that defines *G(n,e).* The list of the current GT descriptors for PNNs and their significance for materials structures is listed in **Table 1**, a spectrum of parameters that can be further expanded as needed. Each of the workflow modules is controlled by settings in the GUI (**Figure S2**) to provide the researcher full access to both the image processing features as well as the GT analysis capabilities.

**Table 1.**
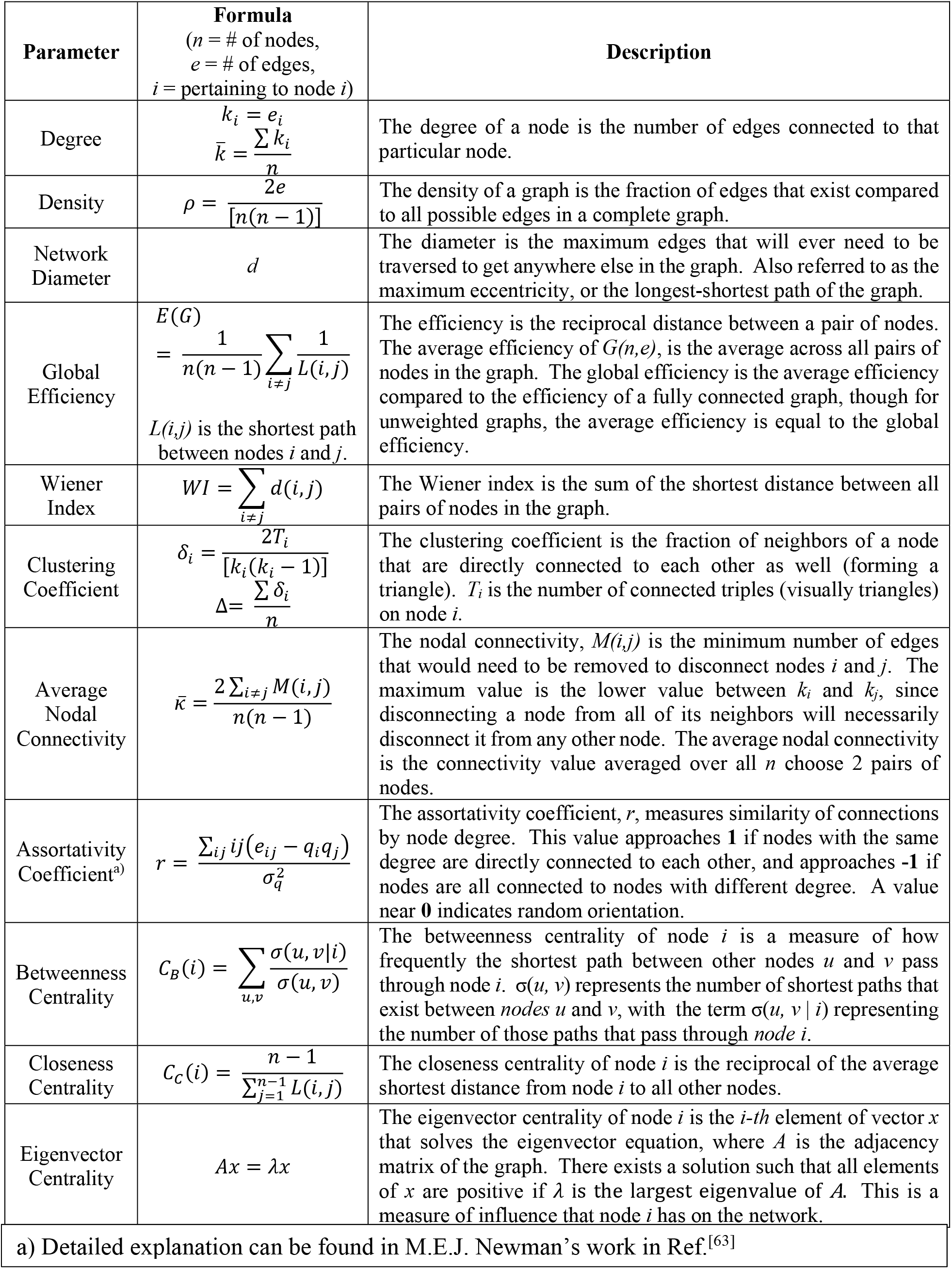
GT parameters for description of percolating network materials.

### 2.1 Image processing

*StructuralGT* initially converts images into an 8-bit gray scale. Using a built-in visualization tool, the user can crop the image to focus on the area of interest, which is necessary for electron micrographs that have scale bars, annotations, or defects. Since every electron microscopy or other image contains noise – even those obtained with noise-reducing hardware – noise reduction and enhancement of the contrast are essential next steps.^[47]^ *StructuralGT* utilizes the median filter and Gaussian blurring filter for noise removal in order to avoid the appearance of the false pixels in the background or, alternatively, their disappearance in the PNN section of the image. The median filter is most suitable for high contrast images with a low amount of noise, being particularly efficient in removing “salt and pepper” noise. While it may struggle with large pixel-to-pixel intensity deviations, this filter is advantageous for *G(n,e)* extraction because it preserves edges, a feature that is key for the conversion of these images to skeletal maps.^[48]^ Alternatively, the Gaussian blur filter has a variable kernel size that allows it to be used on a much larger scale within the image. The blurring can cause degradation of the sharpness of the edges, but it preserves connected features in the network that appear disconnected due to noise or poor contrast. A combination of a median with a Gaussian blur filter **(Figure S3**, top row) is effective for eliminating noise while retaining the connectivity between the junction points.

Images with poor contrast can also be processed by *StructuralGT* taking advantage of the gamma adjustment slider in the GUI to correct non-linear brightness that can aid in enhancing the contrast between features and the background. This slider can be set to any value from 0.01-5.00 (resolution 0.01), with gamma values greater than 1.00 making the image brighter, and gamma values less than 1.00 will make the image darker. Since this is a non-linear correction, pixels with 8-bit values nearer the middle of the 0-255 range will be most affected, while those pixels that are further towards the bright or dark ends of the spectrum are less affected.

For many images, these tools, combined with proper thresholding will be sufficient; however, more complex or noisier images may require some additional processing. *StructuralGT* implements four types of edge detection techniques to assist in specific feature detection. In the GUI, the user can select a Sobel, Scharr, and Laplacian gradient filter included in *scikit-image* module,^[49]^ and an autolevel filter included in the *OpenCV* module.^[50]^ A visual comparison of these methods for enhancing feature detection can be found in the supporting information **(Figure S3)**.

Conversion to a binary image is the final step of image processing when pixels are binned into two categories: 1 for the network and 0 for the background. Three thresholding options are currently used to produce the binary image: global, OTSU, and adaptive. The global threshold option plainly segments the image based on comparing each pixel’s 8-bit value to a user-set value, and the binary assignment is based on whether the pixel is greater than or less than the set threshold. The OTSU thresholding is very similar to the global, with the distinction that the threshold value is not determined by the user but is rather calculated based on Otsu’s binarization to minimize the variance in the histogram above and below the threshold.^[51,52]^ The third option is a Gaussian adaptive threshold where the threshold value is a Gaussian weighted sum of neighboring values. The user supplies a kernel size for the adaptive threshold that defines the size of the neighborhood from which the weighted sum is taken. This effectively applies a threshold in local areas rather than the image as a whole, so that regions in the image with varying brightness or contrast can more accurately apply the threshold. Images with large background regions may have incorrectly attributed network regions, due to a lack of local contrast in these areas, and variable thresholding filter provides a possibility to adapt to these image deficiencies. The GUI displays the current filtered binary image so that the user can quickly identify the optimal conditions for image detection.

These methods can be applied to any material or image type, though certain methods may prove more effective for different samples and types of PNNs. The three thresholding methods display different effectiveness when applied to SEM, TEM, and STEM images **(Figure 2)**. The global threshold excels in instances when there is high and clear contrast, such as in STEM images that are largely planar. However, SEM images representing 3D PNNs usually require one to segment the network in the foreground from lower-brightness network features in the background (**cf**. **Figure 2a**). The OTSU threshold in this case provides a more consistent analysis of PNNs than a unique global threshold. TEM images frequently have a noisy background, and the distribution of pixel values often fall into a singular peak that is difficult to accurately segment without the adaptive threshold. All thresholds can result in disconnected scatter noise that is incorrectly assigned to the network in the binary images, especially for images with low contrast **(cf**. **Figures 2f-h)**. These misidentifications can be corrected by removing disconnected segments during the *Graph Extraction* segment of the workflow.

**Figure 2.**
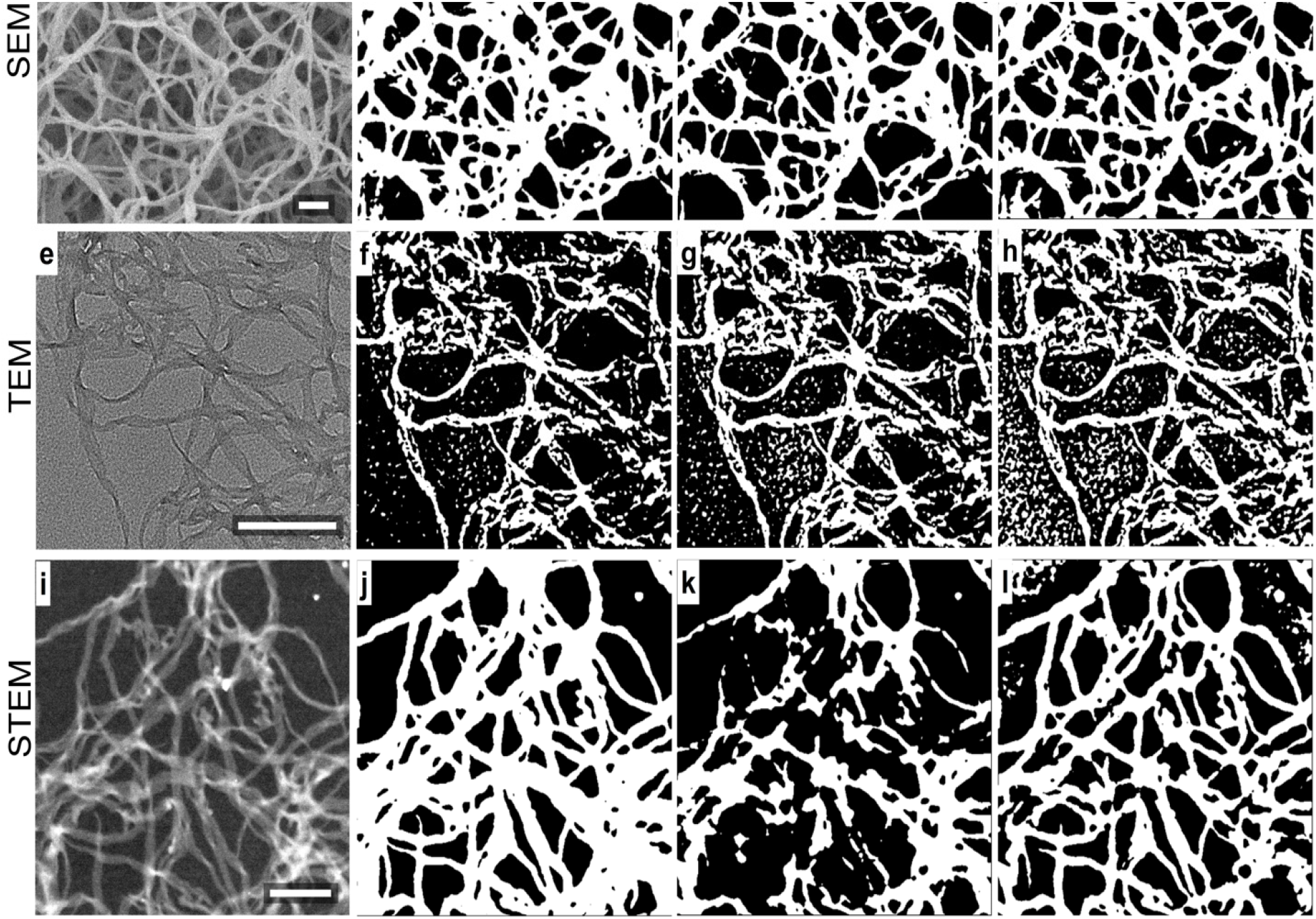
**a-d)** SEM image of an ANF aerogel, and the binary images using different thresholding methods. The global threshold value was 116, and the adaptive threshold kernel size was 155; **e-h)** TEM image of dispersed ANF fibers, and the binary images using each thresholding method. The global threshold value was 108, and the adaptive threshold kernel size was 155; **i-l)** STEM image of dispersed ANF fibers, and the binary images using each thresholding method. The global threshold value was 74, and the adaptive threshold kernel size was 155. A Gaussian blur of kernel size 11 was applied before each threshold. Scale bars are all 200 nm.

### 2.2. Graph extraction

Binary images of networks obtained after thresholding from Section 2.1 are converted into a *G(n, e)* by skeletonization, an image processing step that identifies boundary pixels and removes them in successive passes over the digitized image. The algorithm does not remove the pixels if it would disconnect the PNN fibrils, leaving a 1-pixel wide representation of the features from the binary image. This approach consistently identifies the branches and intersections, a feat that cannot be repeatedly accomplished by manual processing of SEM or TEM images of PNNs and other nanoscale structures.

The resulting ‘*network skeleton*’ is visually similar to a graph but the resulting *G(n, e)* must be a properly-formatted dictionary object for graph algorithms to properly assign edges and nodes to filaments and intersections, respectively. Nodes are identified by scanning through the binary image and classifying each pixel based on how many neighbors it has in the 3 × 3 cell centered around the pixel. Pixels with greater than two neighbors are considered to be branch points, pixels with fewer than two neighbors are considered to be the nodes in the ends of edges, and pixels with exactly two neighbors are considered to be part of a branch. Both end points and branch points are identified as nodes, assigned a numerical identifier, and added to the graph dictionary object. Edges are then identified by tracing the branches that directly connect two nodes, and the edges are added to the dictionary as a tuple of the nodes that the edge connects. Because structural materials are by nature bi-directional (a fiber cannot connect from node *i* to node *j* without also connecting the nodes in the reverse direction), all edges added to the graph dictionary are set to undirected edges in *StructuralGT*. The graph that exists within the ‘skeleton’ image is identified, with end points displayed in red, branch points displayed in blue, and the black pixels representing edges **(Figure 3a)**.

**Figure 3.**
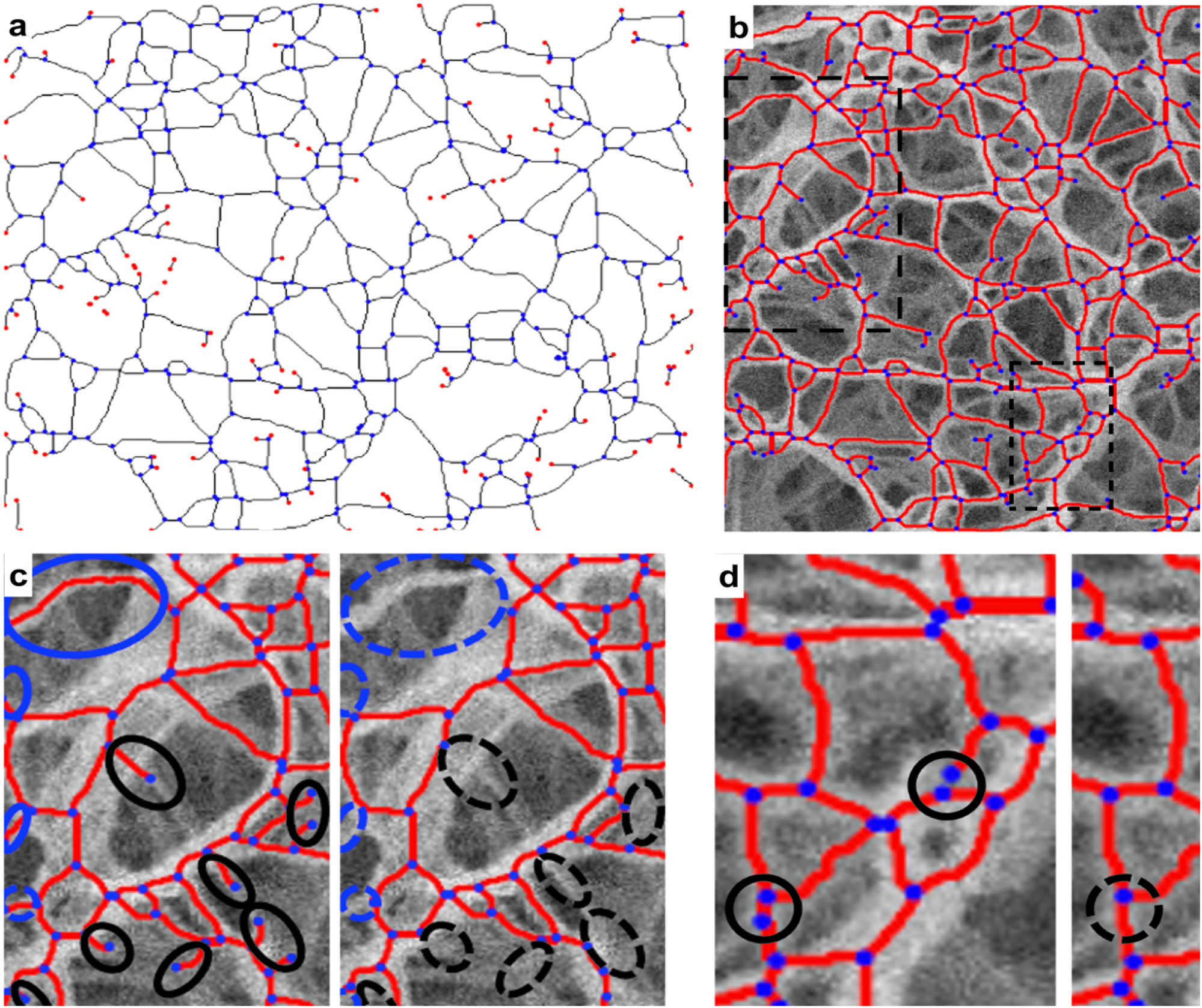
**a)** The ‘skeleton’ image extracted from the binary version of an SEM image of an ANF PNN in **Figure 2c**. Branch points are identified with blue dots, and end points are identified with red dots; **b)** the graph obtained from the skeleton in **3a**, which shows that the disconnected segments are excluded. Nodes are identified with blue dots, and edges are identified with red lines. The long-dashed box indicates the approximate regions of focus for **3c** and the short-dashed box shows the focused region for **3d**; **c) Left:** a zoomed in region from **3b**. **Right:** the same region shown with pruned edges. The ovals highlight affected regions. Black ovals indicate jagged edges that originate from the thresholding, and blue ovals indicate dangling edges created by a fiber getting cutoff at the edge of the image; **d) Left:** a zoomed in region from **b**. **Right:** the same region shown after applying *merge nearby nodes* setting. The circles highlight affected regions. The geometry of the skeletal image will prevent the merging of nodes in some instances, so not all closely located nodes are merged.

Before converting these one-pixel wide images into *G(n,e)* representations of the materials, corrections may optionally be applied in order to improve the accuracy of the ‘skeleton’. These errors can come from several sources. A primary source is from disconnected segments that appear as a result of noise or a difference in contrast that was picked up during the binary segmentation and translated to the skeletal image. These can be erroneous features in the background (**cf**. **Figure 2f-h**) as well as true features that are correctly disconnected. In the case of analyzing a structural network, only the largest connected subgraph would typically be of interest, and several GT parameters (e.g. connectivity) cannot be evaluated if multiple subgraphs exist. Using the option in GUI to *remove disconnected segments* is strongly recommended so that the material representation by *G(n, e)* is a singular subgraph **(Figure 3b)**.

Further corrections to improve the accuracy of the *G(n, e)* representation can be implemented easily by a checkbox in the GUI. Jagged edges (artifacts resulting from imperfect thresholding) and dangling edges (features cutoff by the edge of the image) can be corrected for using a method called pruning **(Figure 3c)**. Correcting for these elements increases the accuracy in many instances. Dangling edges can, for example, distort the calculated parameters (see 2.3) describing the connectivity of the graphs compared to the true values that characterize a near infinite network of the nanofibers or similar filaments since the nodes at the edge of the image will have a connectivity of one instead of their true value. The conversion of the skeleton to a *G(n, e)* representation can also occasionally lead to an over-expression of nodes. Since the nano- or micro-scale fibers in the binary image have an associated width, the intersection of two or more of these fibers can occur over several pixels instead of just at a singular point. The result of skeletonizing an X-junction can then sometimes result in two Y-junction nodes instead of the expected singular node with degree 4. The GUI provides an option that attempts to correct these instances by superimposing a small disk element over the nodes, and repeating the skeletonization, with the *merge nearby nodes* option **(Figure 3d)**.

By default, graphs generated by *StructuralGT* do not allow parallel edges connecting two adjacent nodes (aka multigraphs) because the probability of two or more distinct filaments connecting neighboring nodes without them either collapsing into one or forming additional nodes is low in realistic PNNs. Users, nevertheless, can choose to change these rules for parallel edges and self-loops to fit the needs of the analysis, keeping in mind that several *NetworkX* subroutines calculating betweenness centrality, clustering coefficient, and max flow do not support the calculation on multigraphs.

Weighted network analysis can also be performed by *StructuralGT*. In this case, an additional attribute, i.e. *weight*, is associated with each edge or node. Within the framework of the current package, the weights of edges, *w_e_*, is assigned by calculating the widths of the segmented fibers which is useful for correlations between GT parameters and mechanical properties of the network materials. The trace along each branch can be used to store data on both the pixel width and length of the branch. The length is taken from the skeletal image, whereas the width is approximated from the width of the fiber in the binary image, measured along the perpendicular bisector of the edge’s trace. As with the rest of GT analysis, it is important to remember these measurements are scale-free, so analysis of weighted graphs should be only considered with images on the same physical scale and instrument magnification.

### 2.3. GT parameter calculation

Besides the main result, which is the *G(n,e)* description of the structure of a PNN and similar materials, multiple graph parameters can be calculated. The GT parameters selected for calculations in the GUI may be reported as a single characteristic value, an average value per graph, a histogram displaying the parameter distribution for individual node/edge, or their ‘heat map’ overlaid onto the original image. The GT parameters currently supported by *StructuralGT* are given in **Table 1**; the details of how the calculations are performed can be found in the *NetworkX* documentation.^[53]^

To provide additional details about and physical sense of the parameters that can be used to describe PNNs, here we discuss some of them that can be particularly informative comparing the structure of one PNN to another one. Additional information to aid in understanding and visualizing the GT parameters can be found in the supporting information (**Figure S1** **and** **Table S1**).

The *degree of a node* (*k_i_*) is the number of edges connected to the specific node. This is the simplest parameter describing how connections are made in the network, specifying in particular whether the nanofibers are branching or intersect at a high order. For a percolating network made from a single material, such as ANF, *k_i_* rarely takes a value of 2 for a node since nodes in such networks represent Y- or X-junctions. *k_i_* = 2 for a particular node can be the result of pruning or removing parallel edges. Note that GT descriptions of other materials, where inorganic nanoparticles (NPs) serve as nodes and edges are formed from the organic interfaces or filaments between them, *k_i_* = 2 can be quite common when describing the formation of NP chains.^54^

The density of a graph is simply a ratio of the number of edges in a graph to the number of edges that are possible if all nodes were directly connected by a single edge. Both *k_i_* and *ρ* can be used to characterize, for instance, molecular crosslinking in polymer networks, and the entanglement of polymer fibers.^[55,56]^ Note that typical micrographs are 2D images and thus, the graphs extracted will typically be planar graphs, meaning edges cannot cross without a node at their intersection. Consequently, *ρ* decreases with an increase of *n* if the morphology is preserved.

Characterize the network’s capacity,^[57]^ the *diameter* (*d*)*, global efficiency* (*E*), and *Wiener Index* (*WI*) relate to how easy or difficult it is to traverse the network or its ability for transport, for instance, data. In case of PNNs, ion, electron and load transport through this material can be described by their values. While the *d* represents the maximum eccentricity (i.e. the longest-shortest-path between nodes), *E* and *WI* are the measures of eccentricity through all pairs of nodes in the network. The global efficiency, *E*, characterizes the efficiency for information, energy, or other measures to be transported throughout the network. This can be used to evaluate mechanisms such as charge transport or stress transference in porous and composite materials. *WI* has been widely used in organic chemistry, and has been correlated to properties such as the boiling point, density, and the van der Waals surface area of a material – properties related to the size of a network material.^[58,59]^

The *clustering coefficient* of a node (*δ_i_*) and of a graph (*Δ*) describe the fraction of neighbors of the node that are also directly connected to each other – effectively the fraction of possible triangles that are in the graph. The *nodal connectivity* of a pair of nodes, (*κ*) is the minimum number of edges that need to be removed from the graph in order to disconnect the two nodes from each other.^[60]^ This value spans the range from one to the smallest *k_i_* characterizing the two nodes in question. For PNNs, this lower limit for nodal connectivity is constrained because there must always exist at least one edge between the pair of nodes forming a material. The upper limit for *κ* is bound by the node in the pair of two nodes that has the lower degree, because despite the numerous pathways that might exist between any two nodes, a node can always be disconnected when all of the edges directly connected to it are removed. According to the Menger’s theorem, the value of *κ* for a pair of nodes is equal to the number of edge-independent paths that exist between the two nodes.^[61]^ The value of average nodal connectivity for the entire graph 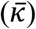 is calculated as the average value for all *n* choosing pairs of nodes in the graph. It is intuitive that both *Δ* and 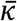 could be related to the mechanical properties of PNN, with the clustering coefficient, *Δ*, being readily related to compression or tensile strength, and the average nodal connectivity, 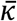, being indicative of resilience, maximum flow, and resistance to shearing.

Different values of *centrality* describe the importance to each node in the graph for the connectivity of the network. The centrality indexes assess and visualize the critical regions of the network, depending on a few different criteria.^[57,62]^ These GT parameters are best evaluated as so-called *heat maps* visualizing the centrality of a given node relative to those around it, rather than as an average value for *G(n,e)* as a whole. The *betweenness centrality* (*C_B_*) describes how frequently a node lies along the shortest path between two other nodes, giving an indication of which pathways are most critical in a network. This analysis can also reveal ‘structural holes’, as well as indicating limitations to the connectivity if there are regions with a large difference in betweenness values. *Closeness centrality* (*C_C_*) shows how close a node is to all other nodes in a graph by finding the shortest path to all other nodes. Identifying large differences in closeness centrality can be useful in determining inefficiencies in a network. *Eigenvector centrality* (*C_E_*) is another metric to evaluate a node’s importance to the network, which is determined by having other nodes with high eigenvector centrality around it.^[62]^ It can help visualize hotspots within the graph; a high value of eigenvector centrality identifies the nodes as being strongly associated to the rest of the network.

The *assortativity coefficient* (*r*) is a measure of similarity. It displays the preference for nodes in a network to be directly connected to other nodes that are the same in some way. Here, the similarity of the nodes is being compared with respect to *k_i_*. Assortativity coefficients range from *r* = −1 for networks where the connected nodes are dissimilar to *r* = +1 for those networks where all nodes are directly connected to nodes that are similar, i.e. their *k_i_* values are the same. Networks with *r* = 0 display a random distribution of similarity for directly connected nodes. This metric is important for demonstrating the presence of local order and heterogeneous segments of *G(n,e)* within a graph. Values of *r* that are very strongly positive or negative indicate ordered homogenous mixing of nodes in the network, whereas moderate values indicate heterogeneity and local areas of similarity in a sea of disorder.^[63]^ When *r* →0, there is no order, though it may be heterogeneously mixed. This GT index could potentially be useful for correlating with different phases in a network, crystallinity in polymers, and hierarchical organization of PNNs. **Figure 4** and accompanying **Table 2** provides a visual comparison and explanation of some of the listed GT parameters, such as *Δ*, 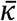, and *r*. Some familiar graphs are compared to an image of ANF to make relationships between GT parameters and network structures more tangible.

**Table 2.**
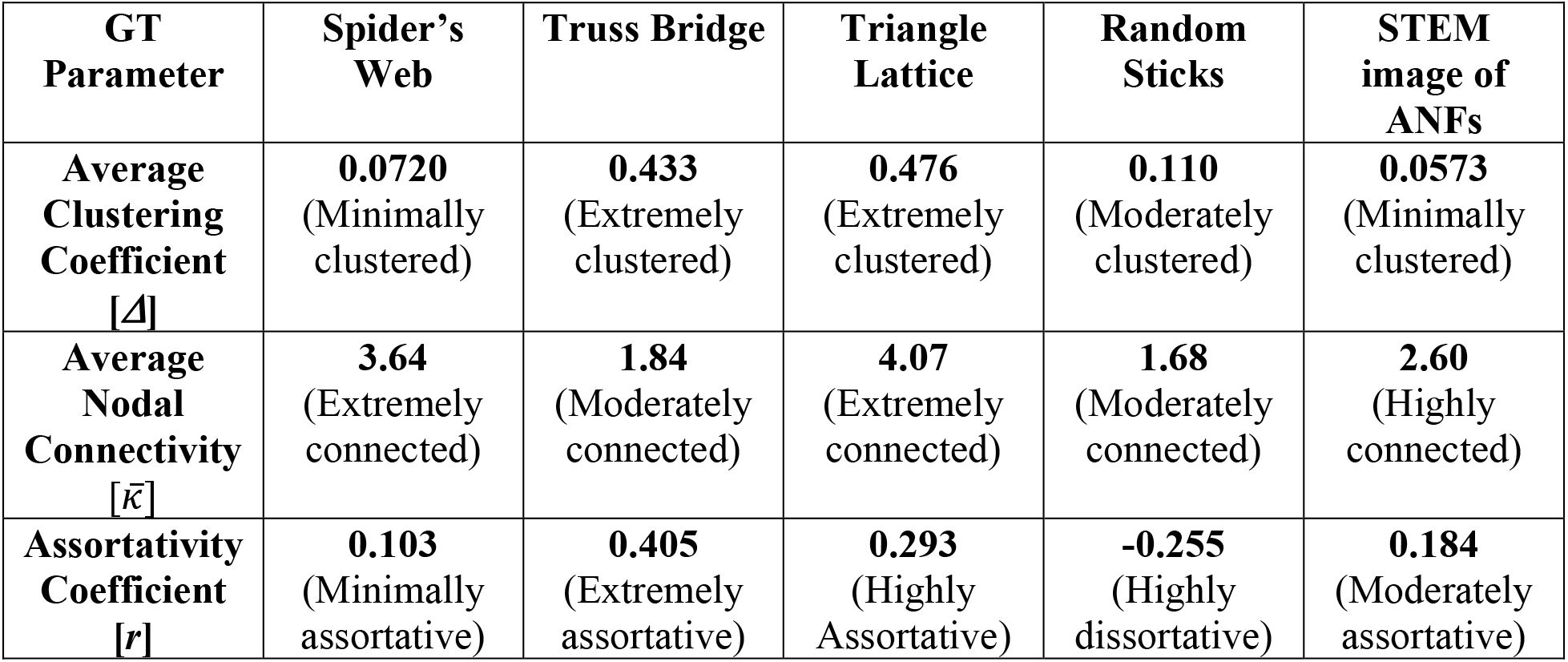
GT parameter values for the graphs in **Figure 4**. The values for average clustering coefficient, average nodal connectivity, and assortativity coefficient are displayed, along with a descriptor to help explain the intensity of the value relative to other structural networks.

**Figure 4.**
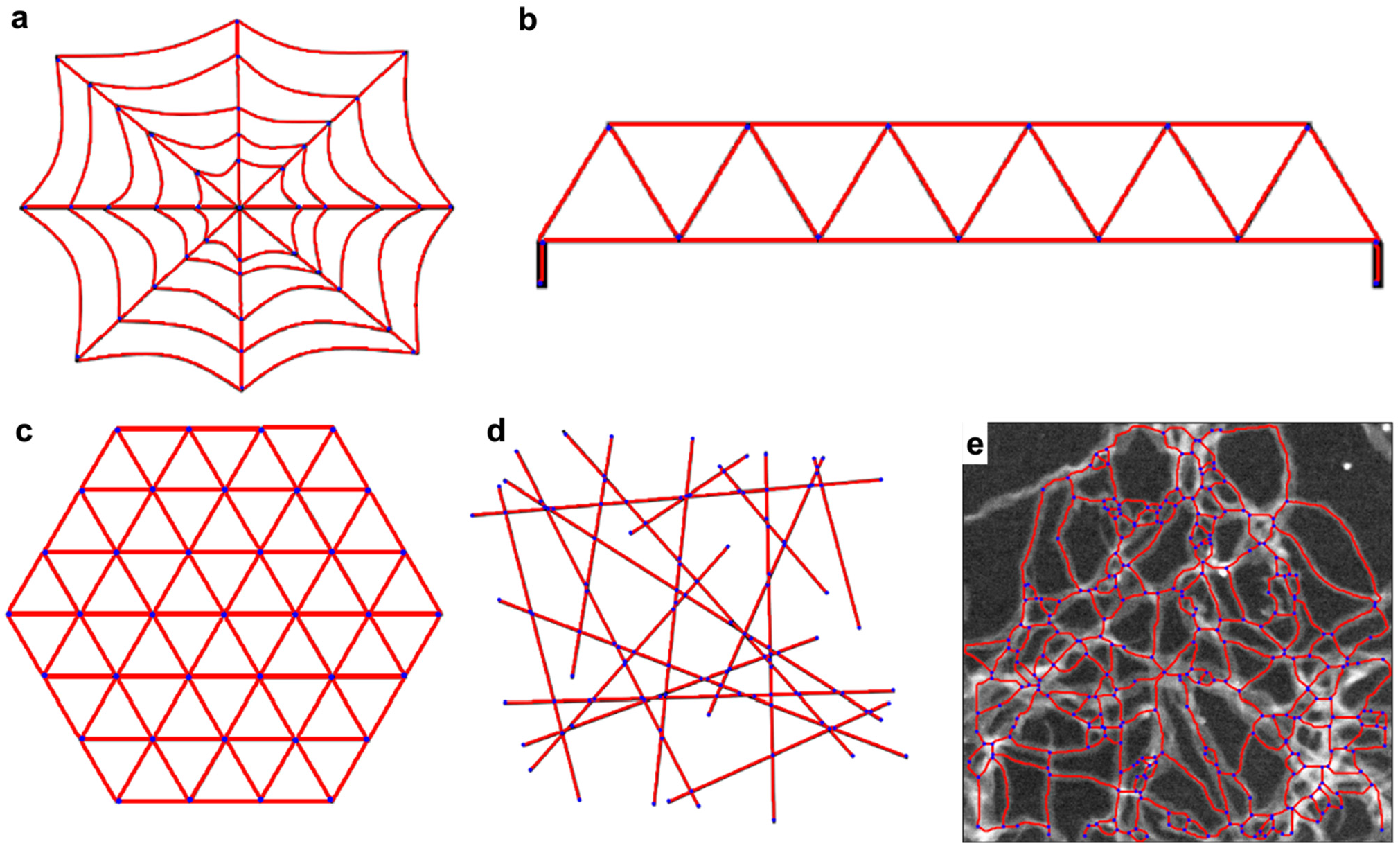
Several types of graphs to better visualize the GT parameters of average clustering coefficient, average nodal connectivity, and assortativity coefficient better. **a)** Schematic illustration of a spider’s web graphic. This structure has a high average nodal connectivity, which can be seen since there exists many independent paths to travel between any two nodes. As such, a spider’s web needs many fibers to be severed to disconnect it. **b)** Illustration of a truss bridge graphic, which are significant for their triangular design. The triangular structure leads to a high average clustering coefficient, since many of a node’s neighbors are also neighbors to each other. Since most nodes have a degree of 4, and are only connected to other nodes with a degree of 4, it is also very assortative. This is in contrast to the spider’s web, where nodes are connected similarly in the polar θ direction, but most are not similarly connected in the radial direction. **c)** Illustration of a triangular lattice. This graph is both very connected and highly clustered, since all nodes are at the corner of triangles, and have degrees of 6 on the interior of the lattice. The *r* is also high, since the nodes on the interior are connected to nodes of similar degree, but this similarity breaks at the boundary of the lattice. The value of *r* would increase as the size of the lattice increases, as the perimeter:area ratio decreases. **d)** Illustrative-schematic of randomly-arranged sticks. This shows contrast to the ordered structures in **a-c**, with a negative value for *r*, which is referred to as being dissortative, meaning most nodes are specifically neighboring nodes of different degree. Assortativity near *zero* reflects a random distribution of nodal degree; assortativity for other parameters can also be computed. The random sticks in (d) happen to display, however, a 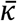 only slightly less than the truss bridge, which is due to the many peripheral edges. e) A graph of a STEM image of ANF, for comparison to the other models.

## 3. Describing the structure of percolating network materials

Parameters, such as the various types of centrality, provide more useful information as a distribution of the nodal values, rather than as an average. Histograms displaying distributions of *k_i_*, *δ_i_*, and the three types of centrality (*C_B_*, *C_C_*, and *C_E_*) provide more detailed information about the GT parameters (and optionally their weighted counterparts) to aid with data analysis of complex networks. Furthermore, these descriptors can be visualized as heat maps where nodes are colored according to their GT parameter and overlaid with the original mage as exemplified for the STEM micrograph for ANF network in **Figure 5a**. In this example, the critical fiber in this structure is easily found by finding the nodes with the highest *C_B_*, whose removal would drastically change the shortest pathways and network connections. This visualization tool provides the opportunity to identify critical features within a complex network that might otherwise be difficult to identify.

**Figure 5.**
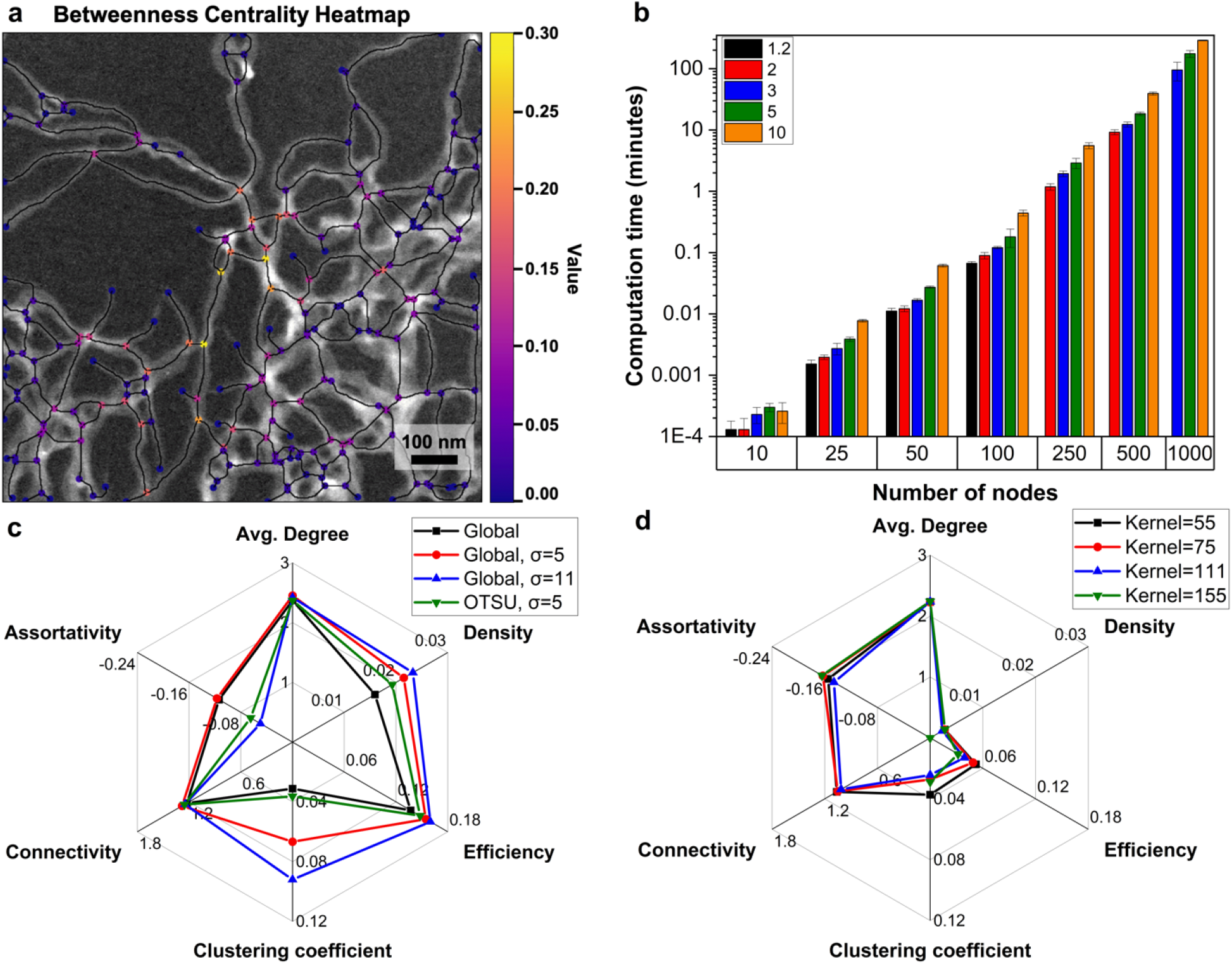
**a)** A heat map displaying the *C_B_* for each node overlaid onto the original STEM image of ANF; **b)** the computation time for calculating the average nodal connectivity for a randomly connected graph with varying numbers of nodes determined by evaluating random graph with a designated number of nodes and edges. The columns are color coded based on the ratio of edges/nodes in the random graph. The number of repetitions - 10; **c)** spider plot describing how several GT parameters of an SEM image of ANF with variations in the Gaussian blur kernel size, σ, and threshold value. The global threshold was value was selected to be 68 by visual appearance of the resulting binary image, and the OTSU determined threshold value was 76. A median filter was applied before processing; **d)** spider plot describing how several GT parameters of a TEM image of ANF change using an adaptive threshold while varying the adaptive threshold kernel size. At a kernel size of 155, the graph became disconnected, so the connectivity could not be calculated.

The GT parameters are robust to the quality of the micrographs and from image to image. Since the GT analysis is performed on data sets contained within a dictionary-of-dictionaries-of-dictionaries object, the obtained GT indexes are identical for repeated analyses of a particular image with identical settings. Comparing the GT analysis of STEM and TEM images of ANF with varying image detection settings reveals robustness of structural analysis by GT **(Figure 5c-d)**. Small variations in the settings cause negligible changes to the reported values of most parameters. The robustness of this image analysis, however, is dependent on the quality of the input image in terms of noise, contrast, illumination, and sample preparation, and more drastic changes (like large changes in the global threshold value) would have a larger impact because the extracted graph would likely become erroneous. The values of *δ_i_* and *Δ* for the STEM image for ANF PNNs varied the most. This sensitivity originates from a reduction in total nodes when the image has small features removed due to applying a larger blurring or local threshold kernel. The binary image in these instances display fewer jagged edges, but at the potential cost of eliminating small true network features. Generally, it is this removal or addition of small features from these small variations in image detection settings that causes changes in parameter values, as the core network structure is constant for well-defined images. When the binary image displays a close match to the resulting graph, then small variations in thresholding, gamma adjustments, or blurring will have negligible effects on the GT parameters.

The calculations are generally computationally quick, although there is an exception. 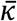 is the only parameter that can take a considerable time for large high-resolution images, particularly if the images possess higher node counts. As the graphs become more complex with a higher number of nodes and higher *e/n* ratio, a significant increase in calculation time is required to calculate 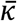 **(Figure 5b)**. This issue arises from having to apply a discrete analysis to determine connectivity that scales with *e*, over *n* choose two pairs of nodes.

## 4. Conclusions

Finding adequate theoretical and computational methods to describe the hidden order of percolating network materials that display a large degree of stochasticity is the key problem addressed in this manuscript. The developed computational tools extract GT parameters from SEM, TEM, and STEM micrographs of a variety of PNNs (**Figure S1**) reporting quantitative descriptors of short and long-range order for seemingly disorganized nanofiber networks. Image processing settings of the *StructuralGT* software package accompanying this manuscript can compensate for image noise, poor contrast, and other defects originating from less-than-ideal imaging conditions. An option for batch processing a folder of many images is also present which would be an asset for ML approaches to engineering of PNNs.

Future developments in this area of materials science include establishing general dependences between GT descriptors and materials properties across different families of PNNs that may be similar to those observed for GT descriptors in chemistry.^[43–46]^ Analysis of 3D networks also represents a potential future development direction of the proposed method of structural description of PNNs.

## 5. Experimental and Computational Methods

### Preparation of ANF samples

Dispersions of aramid nanofibers (ANFs) were prepared following a previously described method by dissolving Kevlar threads in dimethylsulfoxide (DMSO) in the presence of potassium hydroxide.^[7]^ Hydrogels of ANF were produced by spin-coating 2 wt/vol% ANF onto a glass slide and immediately immersing in pure H_2_O. When the water had completely displaced the DMSO in the ANF, the free-standing film was frozen and subsequently dried via lyophilization in order to preserve the network structure for scanning electron microscopy (SEM) imaging. Dispersions of 0.02 wt/vol% ANF were prepared before further dilution with DMSO. Samples for transmission electron microscopy (TEM) and scanning transmission electron microscopy (STEM) images were prepared by drop-casting the dilute ANF dispersion onto the pure carbon film of a copper TEM grid. The samples were dried at 50°C in an oven, under vacuum.

### Electron Microscopy

SEM image analysis was performed on the dried films of ANF networks using a Thermo Fisher Nova 200 Nanolab SEM/FIB. TEM and STEM images were acquired for dispersions of ANF fibers using a Thermo Fisher Talos F200X G2 S/TEM. TEM images were acquired in transmission mode using a Gatan OneView camera, while STEM mode was used to acquire images on the HAADF and DF4 detectors. Images of ANF taken using the DF4 detector were observed to have better contrast than those taken with the HAADF detector. Images were not processed for additional image correction before being read by *StructuralGT* for analysis of the networks in the images.

### Computational Methods

The software’s code is flexible and it can easily incorporate new modules to one of the three core sections: *image detection*, *graph extraction*, and *network calculations*. *StructuralGT* is written in Python version 3.8 and requires the following open-source libraries for computer vision, GT analysis, and plotting of results: *matplotlib,*^[64]^ *networkX,*^[53]^ *numpy,*^[65]^ *opencv-python,*^[50]^ *pandas,*^[66]^ *Pillow,*^[67]^ *scikit-image,*^[49]^ *scipy,*^[68]^ and *sknw*. GT parameters are calculated using *NetworkX module.*^[53]^ Selected images can be cropped and altered to remove noise and enhance contrast to prepare the image for analysis.

For evaluating computational intensity of average nodal connectivity, a brief python script was written in Jupyter Notebook that generated a random graph given a specific *n* and *e* via *NetworkX*.^[53]^ The graph was first checked to make sure it was a connected graph, before timing how long it took to calculate the average nodal connectivity of the graph, and averaged across ten graphs. Lower *e*/*n* ratios were omitted at higher node counts because the fraction of permutations of random graphs that are connected at high *n* and low *e*/*n* represents too small a fraction. The computation was performed on a computer with 8 GB of RAM.

In selecting images for use with *StructuralGT*, it is important to consider several criteria to ensure accurate analysis. The image detection can only identify features through difference in pixel brightness values. Therefore, images that do not form interconnect structures, or are too dense, or have small feature sizes are not recommended. Specifically, networks that are very dense and do not show negative space for contrast will make it difficult to detect the features within the structure, and the program will display connections arbitrarily. Similarly, fibers that are too small (only several pixels wide) may not appear or they may become disconnected when corrections, such as the Gaussian filter, are applied. The magnification should be set so that all features are large enough to be detected accurately. While we have not yet determined what effect varying magnification may have on parameters such as connectivity or clustering coefficient, 2D planar graphs with constant morphology will see that values such as graph density, diameter, and efficiency will scale with the number of nodes in the graph. We recommend that when directly comparing one image to another, that the working distance and magnification be held at a constant value to ensure a quantitatively reliable assessment. Images optimized for high contrast, low noise, and even illumination will obviously produce the best results, and while tools are present to accommodate a wide range of imaging conditions, additional external image correction software may provide solutions to detect challenging images.

A PDF file containing the results of the analysis by *StructuralGT* is saved to the designated directory automatically at the end of the analysis. The resulting PDF displays images of both the initial ‘skeleton’ and the final corrected *G(n, e)* representation (superimposed on the source image) that is used for subsequent analysis of the PNN’s structure. On the skeletal image, the pixels are color-coded as blue for branch points, red for endpoints, and white for part of a branch. On the final graph image, the nodes are labeled as blue dots and the edges as red lines.

The first two pages of the output PDF file are images showing the step-by-step progression from the input image to the final graph. The following page in the PDF shows the selected GT parameters which are displayed in an output table. Some of these parameters are simple while others express averages across all nodes or all pairs of nodes. If the user utilizes edge weights, the weighted and unweighted GT parameters are displayed separately.

*StructuralGT* provides many options for GT analysis, but one may want to explore other processing methods for analysis of *G(n,e)*. Graphs generated from the structural images in *StructuralGT* can be exported as graph exchange XML files (GEXF file, .gexf extension) that can be imported into commonly used GT analysis software, such as the program *Gephi*. The GEFX files store the node, edge, and attribute data while maintaining consistent identifier labels, which allows for the transfer of graphs identified in *StructuralGT* to other platforms, enabling complementary analyses.

## Supporting Information

Supporting Information is available from the Wiley Online Library or from the author.

## Acknowledgements

The primary support of this research is provided by AFOSR FA9550-20-1-0265, Graph Theory Description of Network Materials. The essential support for this study is also provided by Vannevar Bush DoD Fellowship to N.A.K. titled “Engineered Chiral Ceramics” ONR N000141812876, ONR COVID-19 Newton Award “Pathways to Complexity with ‘Imperfect Nanoparticles” HQ00342010033. This work was supported by the US Department of Homeland Security (2015-DN-077-097) and by DARPA (D18AP00063) and DTRA (HDTRA1-20-2-0002) of the US Department of Defense. The authors acknowledge the financial support of the University of Michigan College of Engineering and technical support from the Michigan Center for Materials Characterization. We thank Mark A. Wrobel of DHS and DARPA for helpful discussions during the development of the code for application to sensing materials. We also thank Anirudh Cowlagi of the University of Pennsylvania who assisted with initiating a Python approach to analyzing digital images. This support does not constitute an express or implied endorsement on the part of the government.

## Supporting Information

**Figure S1.**
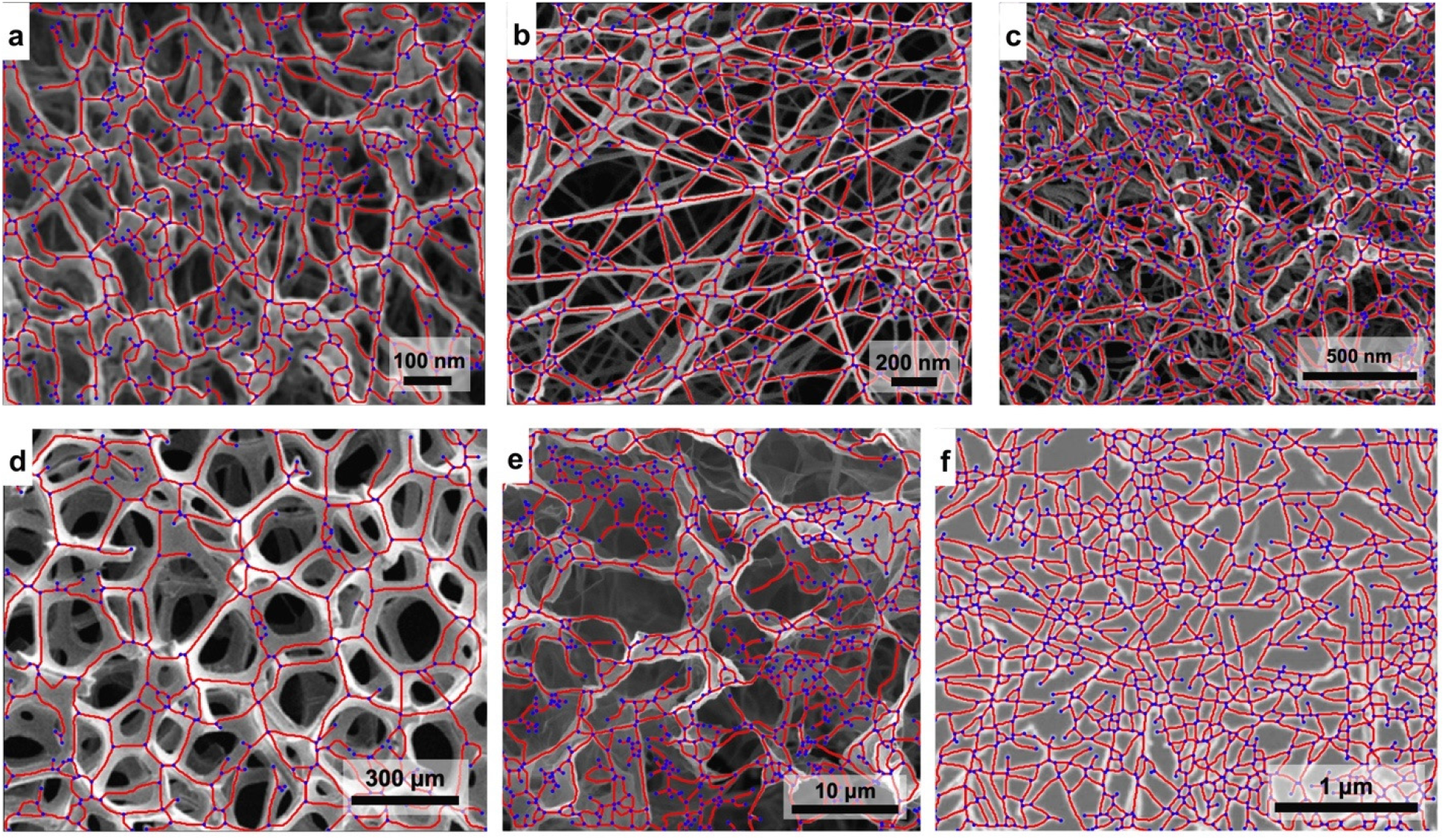
Graphs of the networks displayed in **Figure 1**. a) Graph of articular cartilage collagen; b) Graph of bacterial cellulose; c) Graph of boron nitride nanotubes; d) Graph of nickel foam; e) Graph of graphene oxide and Ca^2+^ hydrogel; f) Graph of single walled carbon nanotube networks.

**Table S1.**
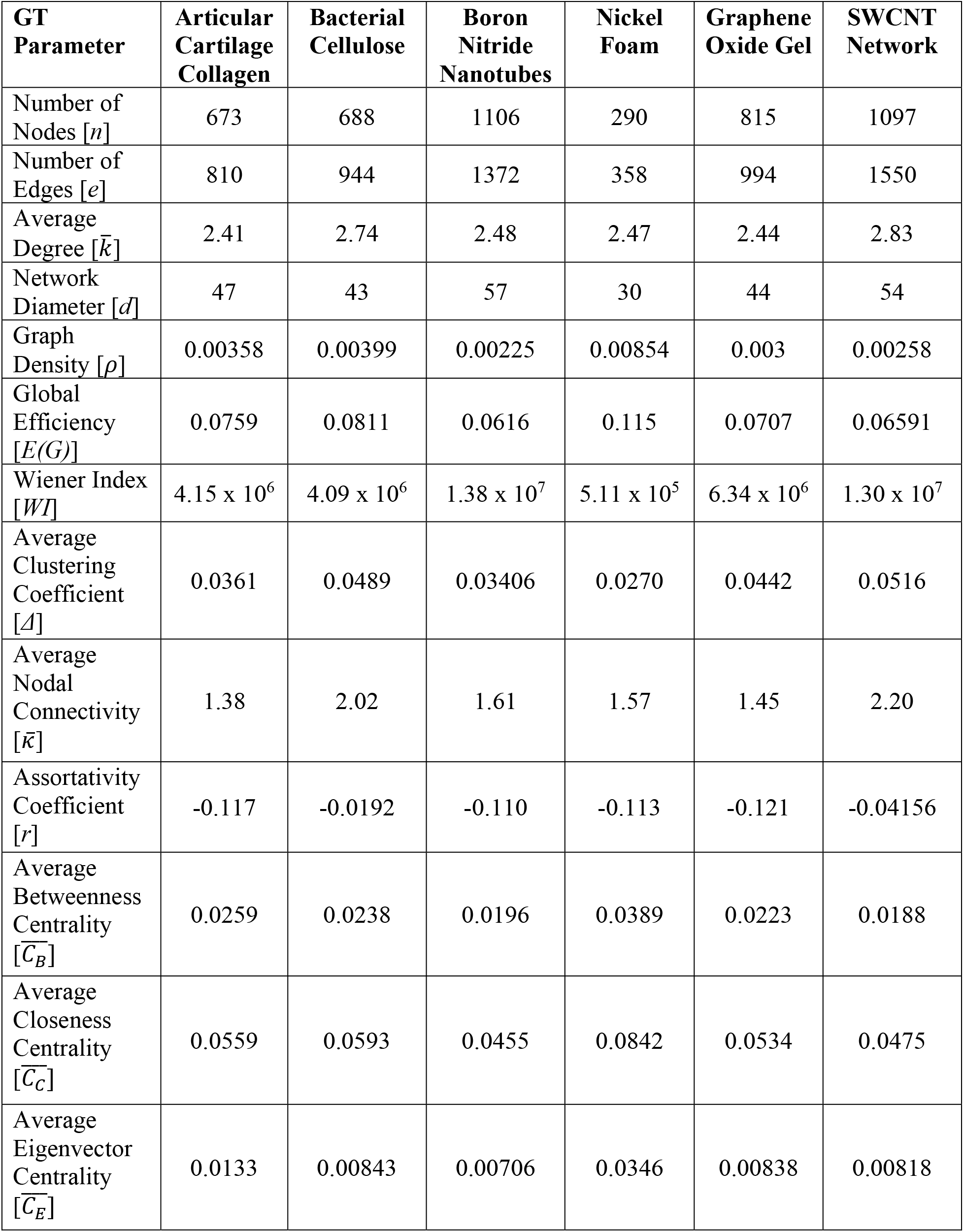
Summary of the GT parameters calculated for the graphs displayed in **Figure S1**.

**Figure S2.**
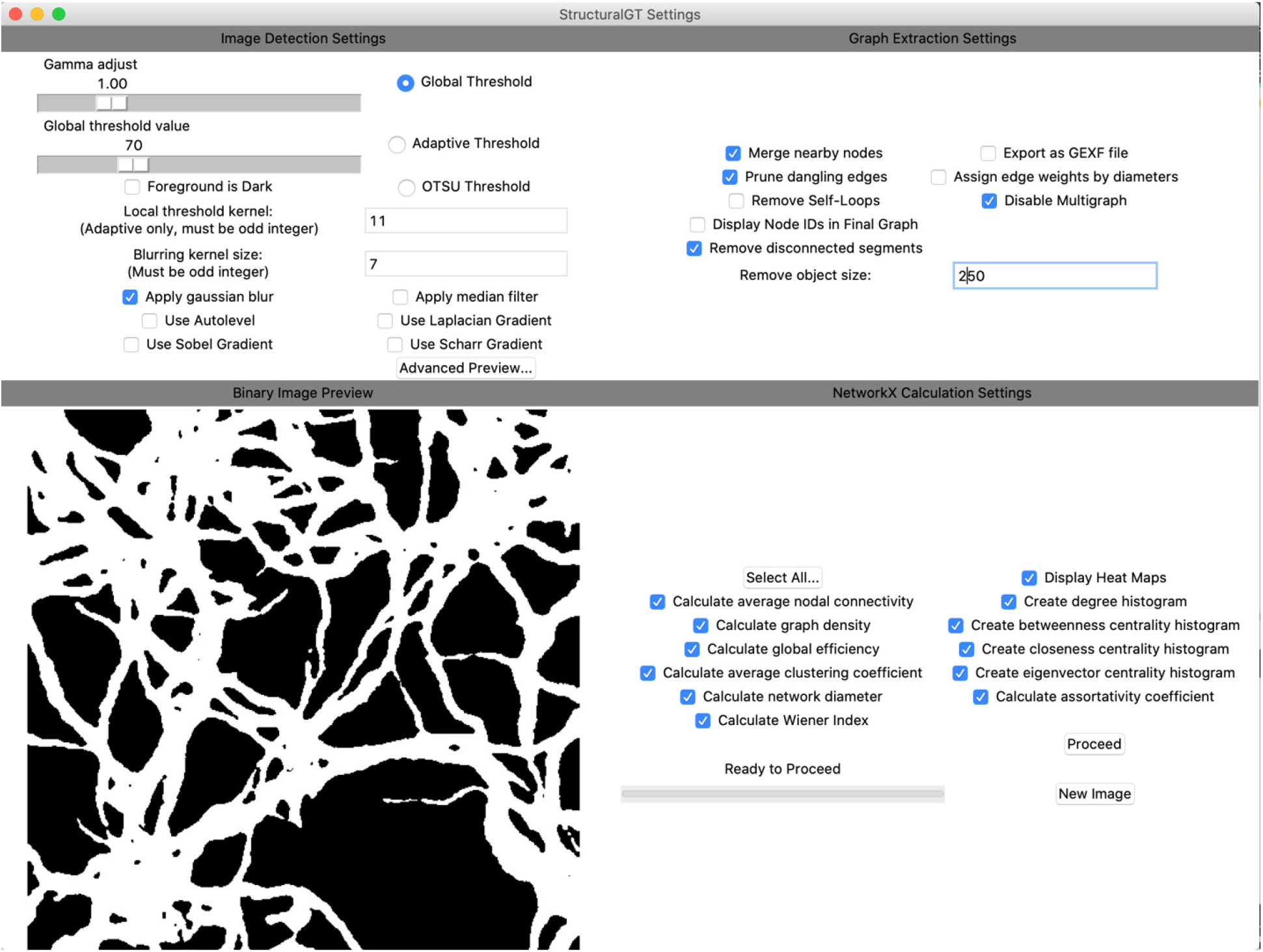
Display of the GUI in *StructuralGT*. Top left shows the settings for performing image detection. Bottom left shows a preview of the binary image that is obtained using the current image detection settings. “Advanced Preview…” can be used to get a more detailed step-by-step preview of the image segmentation. Top right shows the settings for obtaining a graph from the binary image. Bottom right shows the calculations to perform, as well as the instructions how to present the data.

**Figure S3.**
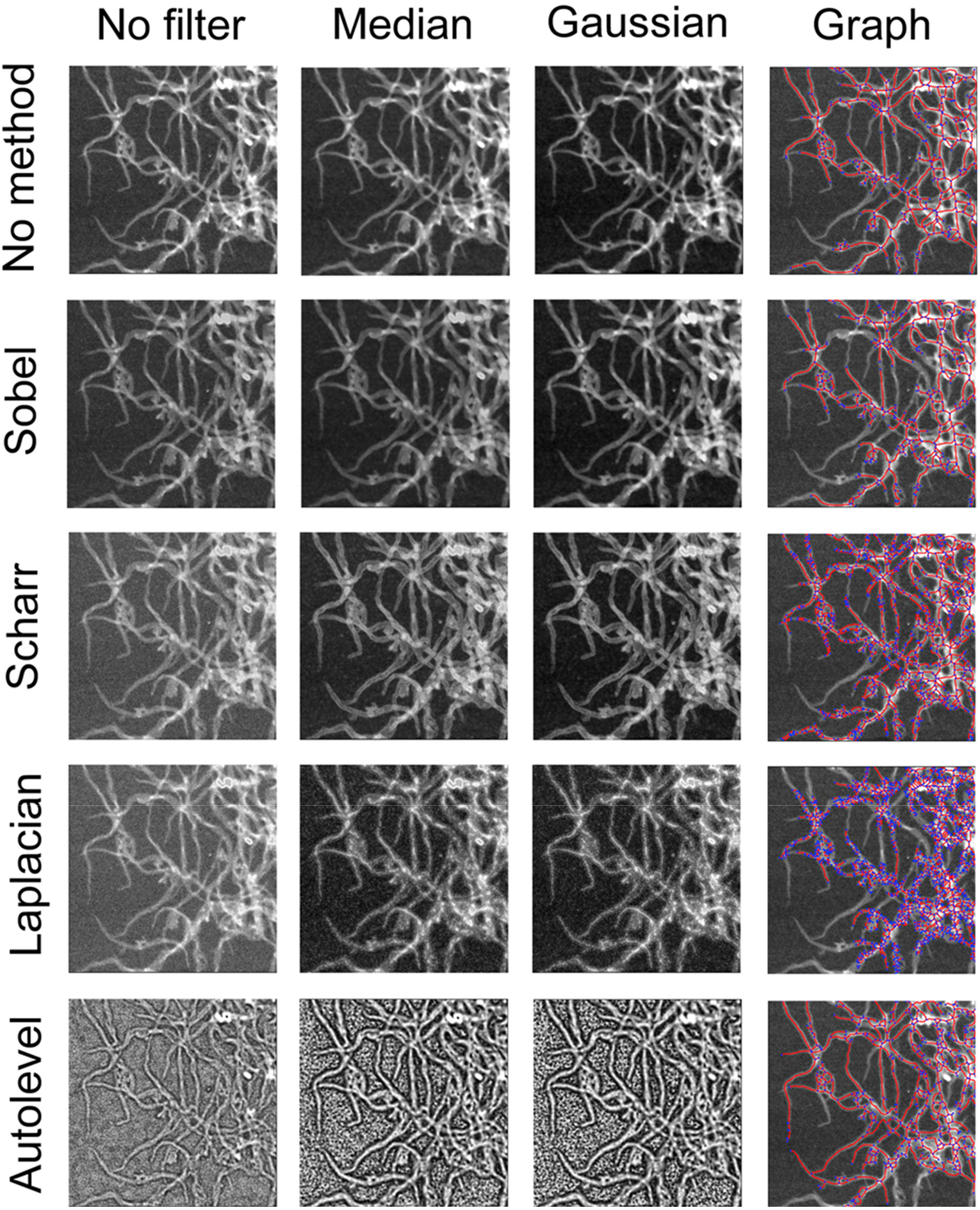
Comparison of several image filtering (in the columns) and edge detection techniques (in the rows) applied to a STEM image of ANF. The Gaussian blur uses a kernel size of 7. The graph obtained for each method was obtained by applying the Gaussian blur before the stated edge detection method, then segmented using an OTSU threshold, and removing disconnected segments. Note that an appropriate global threshold or adaptive threshold may obtain a more accurate graph, but OTSU was used here for consistency in comparison.

The three gradients work methods operate upon the same principle by finding the intensity differences between sections of pixels, and where the image changes greyscale color sharply, the technique identifies these as edges. The Sobel and Scharr gradients operate similarly, detecting edges through brightness gradients, with the Scharr gradient seeking variations in all directions, as opposed to Sobel method that only searches along the vertical or horizontal axes of the image. The Laplacian gradient uses a Laplacian operator to detect edges. This map of edges is then overlaid with the original micrograph to “sharpen” the outline of the image, a process that works well with images that possess low noise but faint details, as these gradients can pull the edges forward to be picked up by the threshold. While reducing blur, the Sobel and Scharr gradients edge detection algorithms have limited use, however, because they may also create false double connections between segments. Of these three gradients, the Laplacian filter tends to bring out the edges the strongest **(Figure S3**, **row 4)**.

Additionally, the image can be autoleveled. This image processing procedure takes a histogram of all pixel values and stretches it to cover the full range of greyscale, effectively scaling the grayscale pixel values from 0 to 255. With low contrast images, autoleveling can be particularly effective, as it increases the contrast between the lightest and darkest pixels. However, the same operation may also bring a large amount of noise **(Figure 2**) and the user will need to rely on proper thresholding and graph extraction to acquire an accurate *G(n,e)* representation of the PNN.

